# Seasonal influenza circulation patterns and projections for Sep 2017 to Sep 2018

**DOI:** 10.1101/191676

**Authors:** Trevor Bedford, Richard A. Neher

## Abstract

This report details current seasonal influenza circulation patterns as of Sep 2017 and makes projections up to Sep 2018 to coincide with selection of the 2018 Southern Hemisphere vaccine strain. This is not meant as a comprehensive report, but is instead intended as particular observations that we’ve made that may be of relevance. Please also note that observed patterns reflect the GISAID database and may not be entirely representative of underlying dynamics. All analyses are based on the nextflu pipeline [1] with continual updates posted to nextflu.org.

**A/H3N2:** H3N2 continues to diversify with many coexisting clades, all of which carry several amino acid mutations at previously characterized epitope sites. The majority of viruses fall into the 3c2.a clade which has been dominating globally for *>*3 years, but 3c3.a viruses continue to persist. The common ancestor of circulating H3N2 viruses is now more than 5 years old, which is rare for H3N2. Despite extensive genetic diversity, serological assays suggest limited, but non-zero, antigenic evolution. We expect multiple competing clades within 3c2.a to persist into the future with no clear immediate winner.

**A/H1N1pdm:** A clade comprising mutations S74R and I295V has recently risen to *>*60% global frequency. Although it shows no antigenic distinction by ferret HI data, the rapidity of its rise suggests a selective origin.

**B/Vic:** A clade with a two amino acid deletion 162-/163-has altered serological properties and is increasing in frequency, albeit slowly. Two other clades (carrying mutations K209N and V87A/I175V) have increased in frequency moderately.

**B/Yam:** A clade comprising M251V within clade 3 viruses continues to dominate. The is little genetic differentiation within this clade and no evidence of antigenic evolution.

## A/H3N2

H3N2 continues to diversify with many coexisting clades, all of which carry several amino acid mutations at previously characterized epitope sites. The majority of viruses fall into the 3c2.a clade which has been dominating globally for >3 years, but 3c3.a viruses continue to persist. The common ancestor of circulating H3N2 viruses is now more than 5 years old, which is rare for H3N2. Despite extensive genetic diversity, serological assays suggest limited, but non-zero, antigenic evolution. We expect multiple competing clades within 3c2.a to persist into the future with no clear immediate winner.

We base our primary analysis on a set of viruses collected between Oct 2015 and Aug 2017, comprising *>*800 viruses per month in Dec 2016 to Mar 2017 and 100-200 viruses per month more recently (Fig. 1). We use all available data when estimating frequencies of mutations and weight samples appropriately by regional population size and relative sampling intensity to arrive at a putatively unbiased global frequency estimate. Phylogenetic analyses are based on a representative sample of about 2000 viruses.

**Figure 1.**
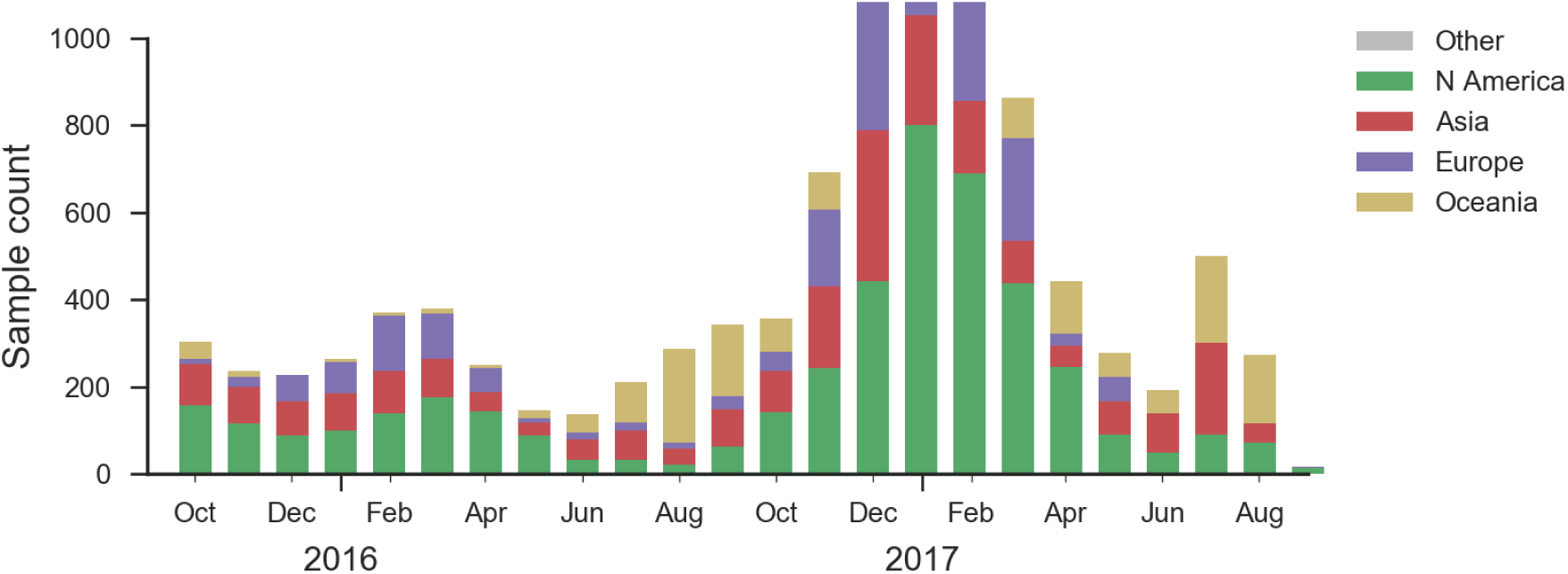
Sample counts through time and across regions. This is a stacked bar plot, so that in all months there are at least 100 samples and in some months there are *>*500 sequences.

At the level of major clades, 3c2.a has continued to dominate while 3c3.a persisted at about 20% frequency in Oceania and North America in the 2016-2017 season (Fig. 2). Within 3c2.a, the clade 3c2.a1 comprising HA1 mutation N171K and HA2 mutations I77V and G155E had risen rapidly in early 2016 but later stabilized at about 50% with somewhat higher frequencies in Europe.

**Figure 2.**
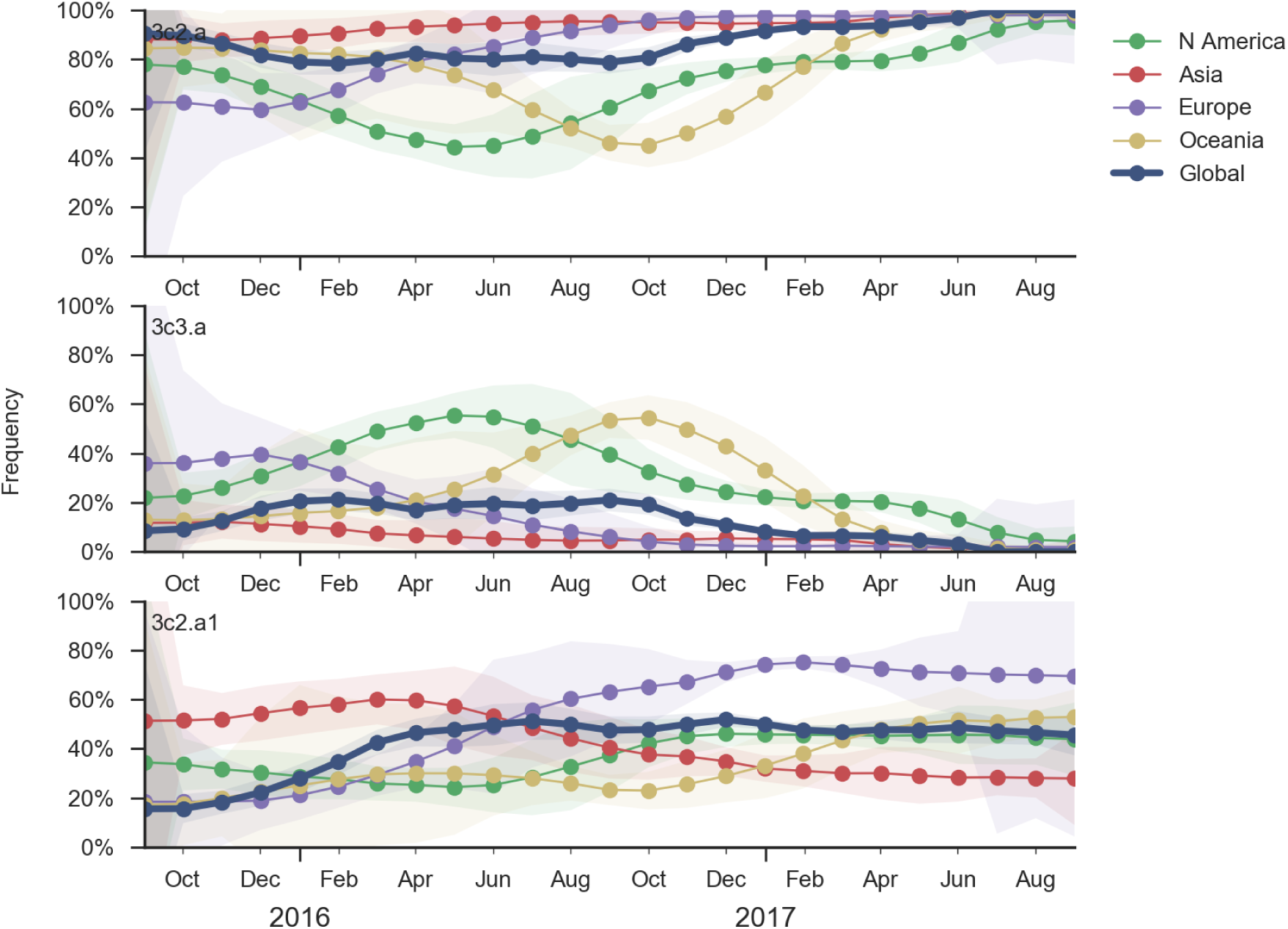
Dynamics of major H3N2 clades. Clade 3c2.a continues to dominate, clade 3c3.a continues to circulate at low frequency, and 3c3.b has not been observed for several months (not shown). Within 3c3.a the subclade 3c2.a1 has persisted at *∼*50% for the previous year. Viruses belonging to 3c2.a but not 3c2.a1 are also at *∼*50% frequency.

We observe substantial diversification within the 3c2.a clade over the past year. There are 5 major subclades within 3c2.a that have reached appreciable frequency (Fig. 3). All of these clades possess mutations at previously characterized epitope sites. These clades are characterized by the following mutations and highlighed in the phylogenetic tree in Fig. 3:

1. Clade 1: 53N, 144R, 171K, 192T, 197H
2. Clade 2: 121K, 144K
3. Clade 3: 131K, 142K, 261Q
4. Clade 4: 135K, HA2:150E
5. Clade 5: 92R, 311Q

**Figure 3.**
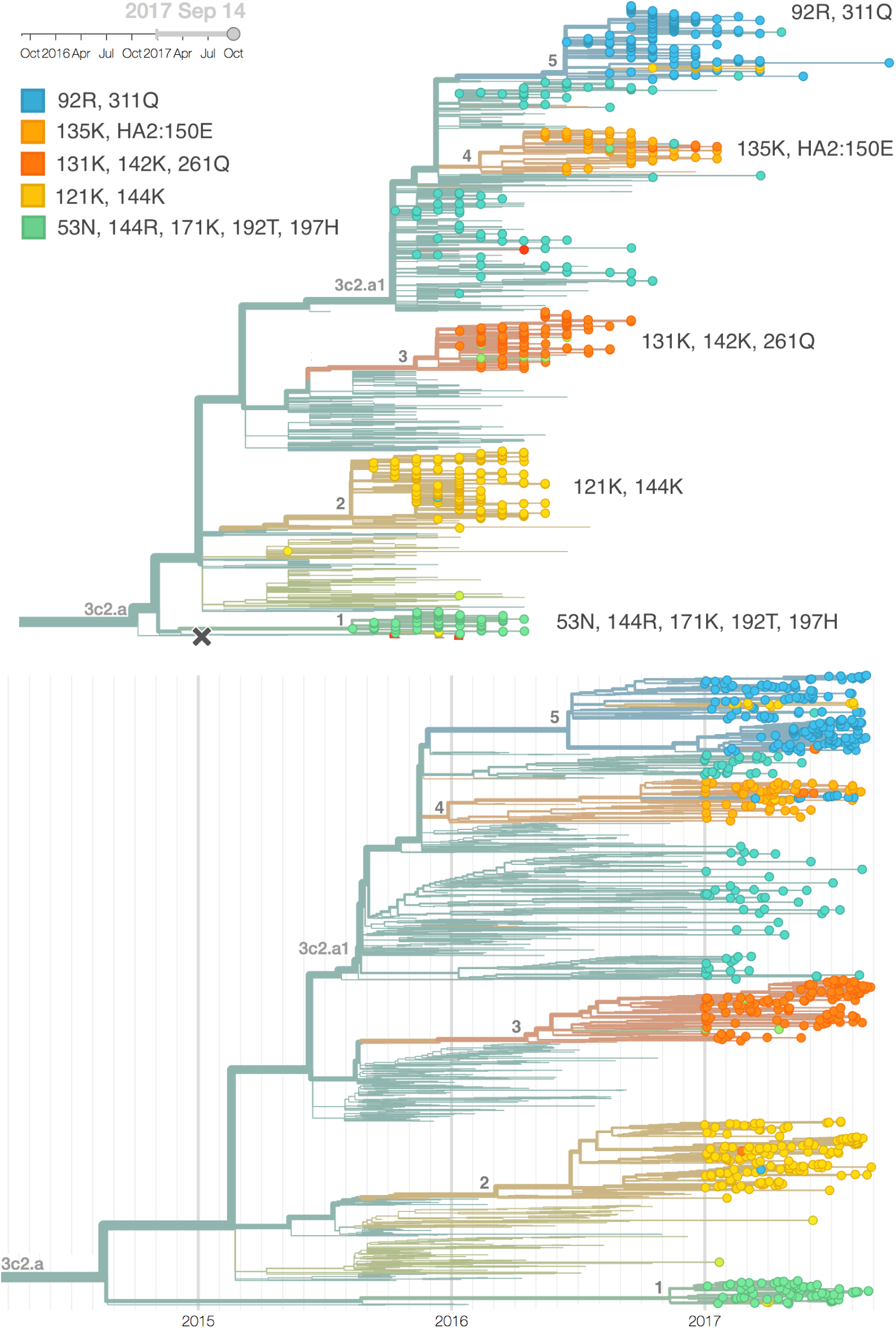
H3N2 / 3c2.a phylogeny colored by genotype. Colors are chosen to highlight clades 1–5. Clades are in reverse order, that is clade 5 is on top, clade 1 is at the bottom. Each of these clades is non-nested and so all compete against each other. Also shown is a time calibrated phylogeny with equivalent coloring.

All of these clades have risen in frequency and are now competing with one another (Fig. 4, Fig. 5). Clade 1 (53N, 144R, 171K, 192T, 197H) is highly derived in terms of sequence, but is at low frequency globally (currently less than 1%) despite making up the a substantial fraction of 2017 Oceania viruses. Clade 2 (121K, 144K) had a large upswing around Dec 2016, but has since declined slightly to ∼ 27% global frequency, but is now increasing again. Clade 3 (131K, 142K, 261Q) has steadily climbed to ∼ 39% global frequency and importantly is at relatively high frequency in Asia. Clade 4 (135K, HA2:150E) has persisted at low frequency and is currently at ∼ 8% globally. Clade 5 (92R, 311Q) has recently increased to ∼ 25% in global frequency, also with relatively high frequency in Asia.

**Figure 4.**
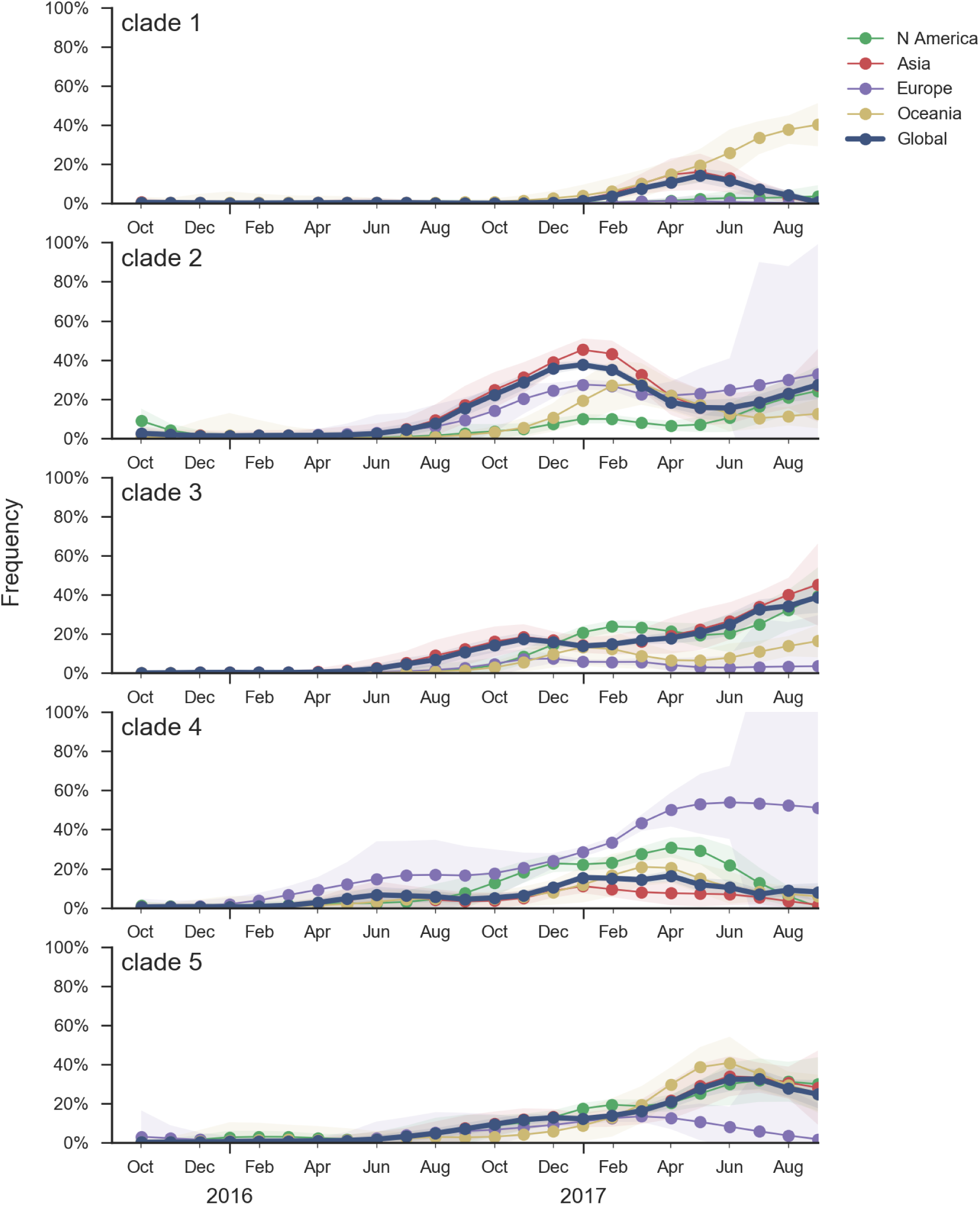
Frequency trajectories of 3c2.a subclades partitioned by clade then by region. We estimate frequencies of different clades based on sample counts and collection dates. Each clade is estimated according to a characteristic mutation unique to this clade. These estimates are based on all available data and global frequencies are weighted according to regional population size and relative sampling intensity. We use a Brownian motion process prior to smooth frequencies from month-to-month. Transparent bands show an estimate the 95% confidence interval based on sample counts.

**Figure 5.**
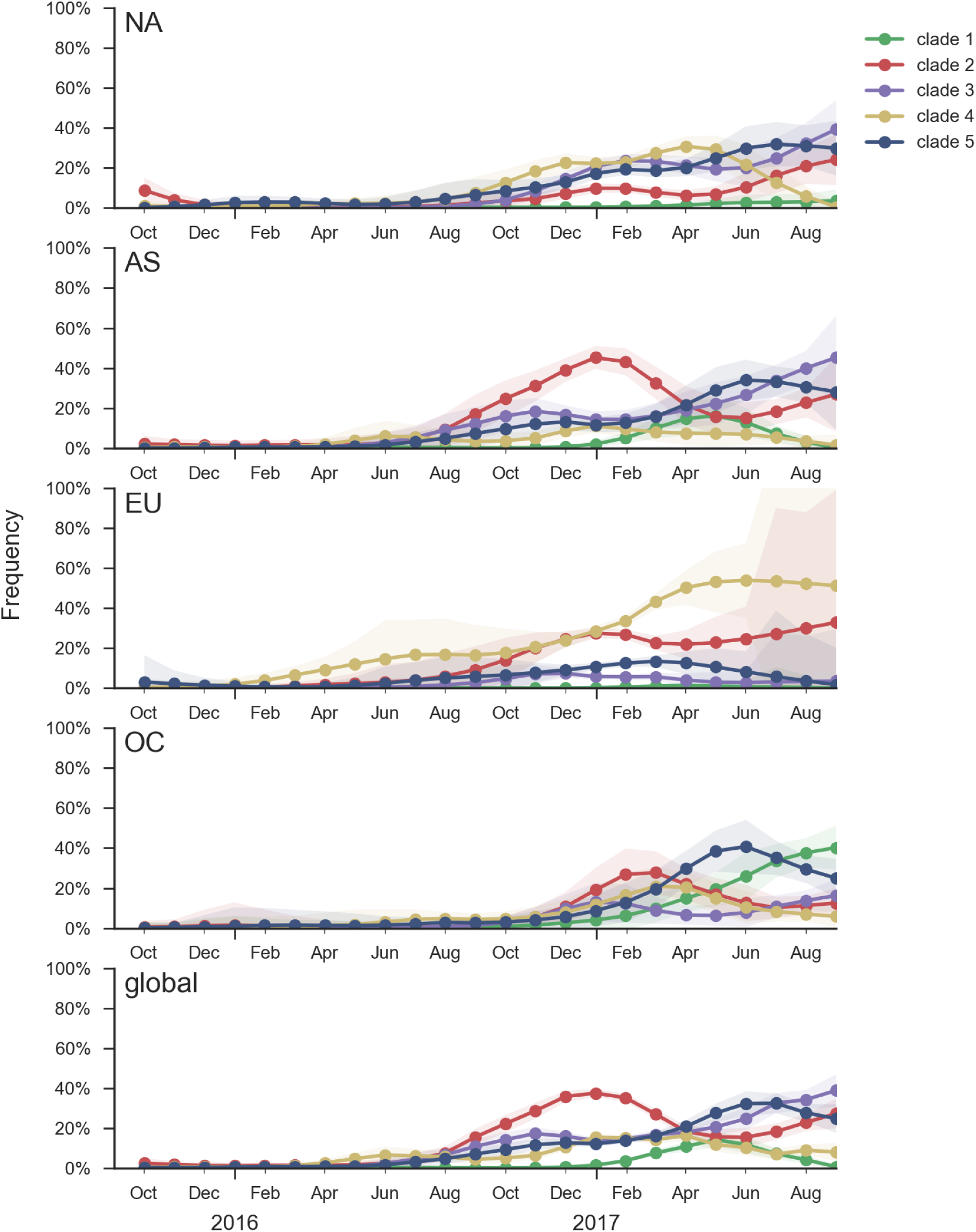
Frequency trajectories of 3c2.a subclades partitioned by region then by clade. We estimate frequencies of different clades based on sample counts and collection dates. Each clade is estimated according to a characteristic mutation unique to this clade. These estimates are based on all available data and global frequencies are weighted according to regional population size and relative sampling intensity. We use a Brownian motion process prior to smooth frequencies from month-to-month. Transparent bands show an estimate the 95% confidence interval based on sample counts.

All 5 of these clades appear somewhat drifted according to antigenic analysis [2] of HI measurements provided by the Influenza Division at the US CDC (Fig. 6). Antigenic differences from vaccine strain A/HongKong/4801/2014 are estimated to be between 0.5 and 2.2 antigenic units (or up to a four-fold HI dilution).

**Figure 6.**
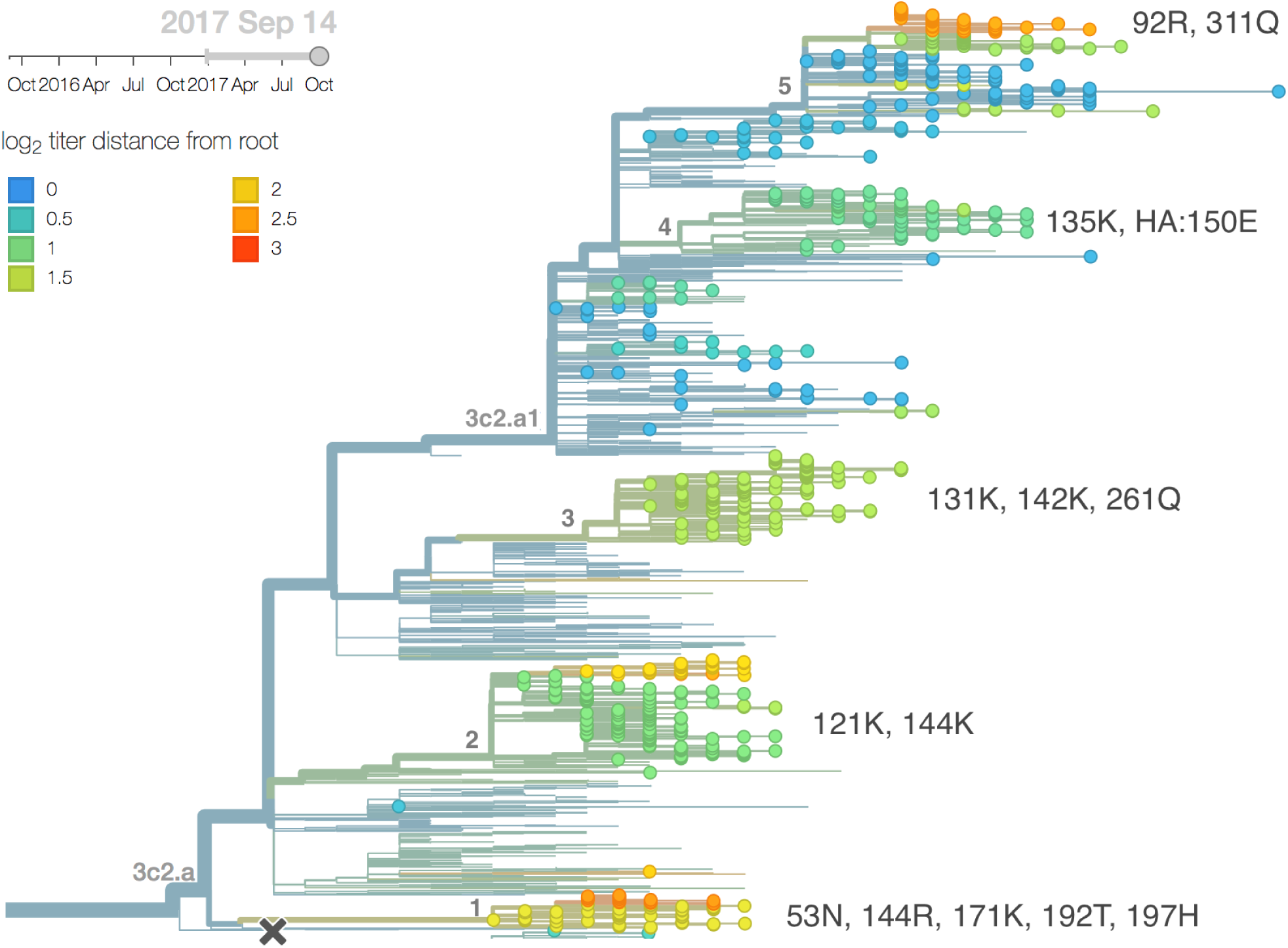
H3N2 phylogeny colored by antigenic advancement. Model estimates of antigenic divergence relative to the root of the tree, which is similar to the titer drop relative to A/HongKong/4801/2014 or A/Michigan/15/2014. Cooler color indicates greater antigenic similarity (less titer drop going from homologous to heterologous titers).

The mutation T135K has appeared independently in clades 2, 4 and 5 and is now globally at 31% frequency. The most rapidly growing clade carries mutations N121K/K92R/H311Q/E62G/R142G/T135K. This is a subclade of clade 5 above with addition of 62G, 142G and 135K. The model fit to the antigenic data estimates that this subclade of clade 5 has drifted by 2.2 antigenic units, 1 of which is derived from the E62G/R142G mutations and 1.2 of which is derived from the T135K mutation (on this background), see Fig. 7. Note, however, that there is no evidence of antigenic evolution in the FRA data (one data point only). A small sister clade with mutation T135N (also part of clade 5) recently developed. Taken together, changes at position 135 likely have selective origin. Previously, the mutation K135T was associated with the Sidney/97 antigenic transition as an “accessory” mutation [3].

**Figure 7.**
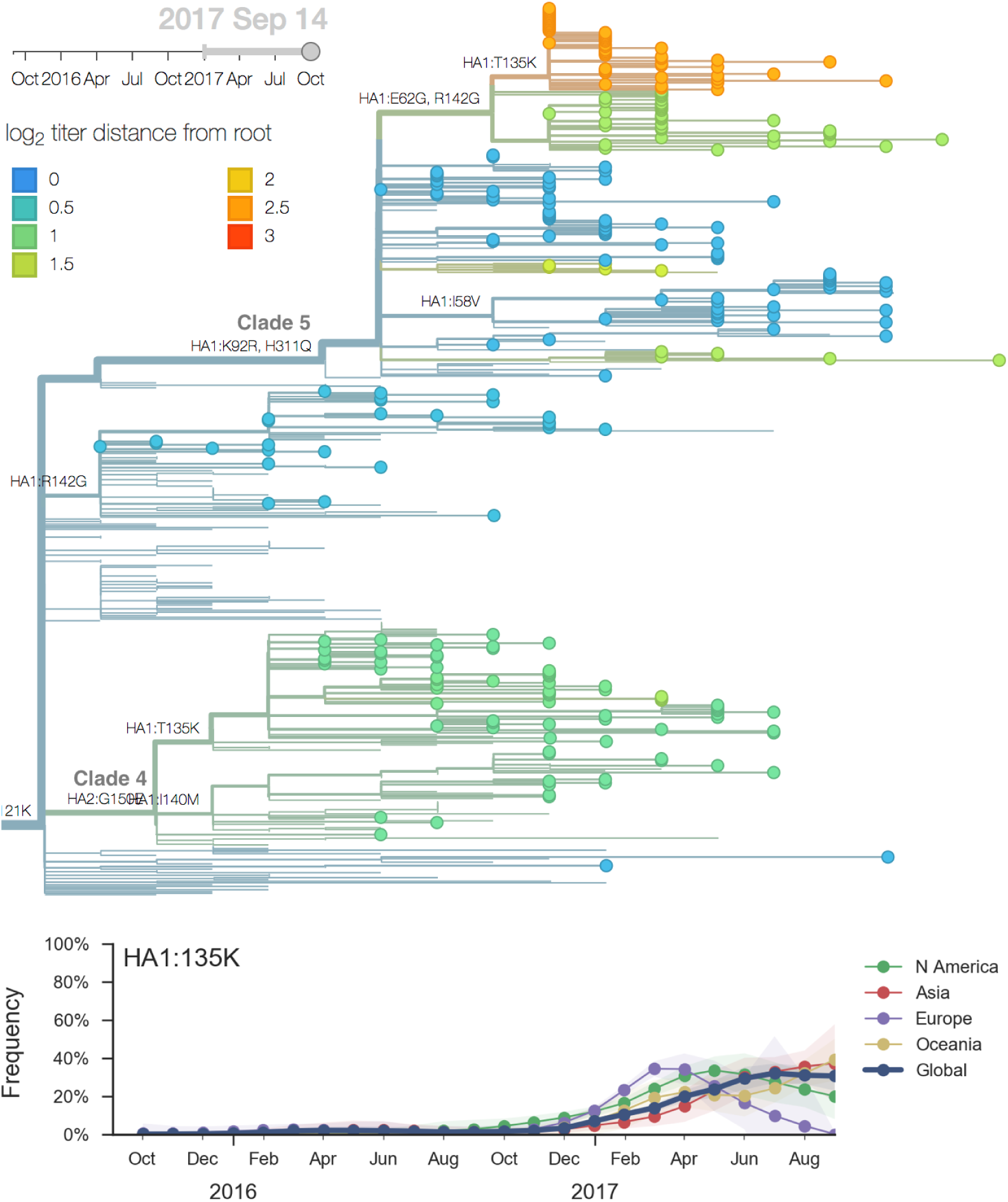
Zoom into subclade of 3c2.a1 characterized by mutation 121K comprising clades 4 and 5, colored by antigenic advance. A subclade within clade 5 bearing a T135K mutation has recently risen in frequency and has drifted *>*2 antigenic units. Parallel T135K mutations in clades 4 and 5 combine together to show an rapid recent increase in frequency.

Clades 1, 2 and 3 are part of 3c2.a but outside of the 3c2.a1 subclade.

Clade 1 with substitutions N31S/D53N/S144R/N171K/I192T/Q197H was observed globally over the past 9 months but is now decreasing everywhere except Oceania. HI data and the model fit suggest a moderate (4-fold, about 2 antigenic units) titer drop relative to A/HongKong/4801/2014.

Despite the titer drop, this clade seems to be outcompeted almost everywhere (with the exception of Oceania). It seems likely that this clade will die back with the ending of the Oceania influenza season.

Clade 3 with T131K/R142K mutations continues to be common in China (but not in South Asia) with an embedded cluster of South and North American isolates from last summer. Subclades within this clade have been expanding recently, and there is evidence for a small amount of antigenic evolution in the HI (0.9 antigenic units) and FRA data (0.1 antigenic units). The 131K substitution is not seen outside this clade.

The most notable subclade of 3c3.a is a clade carrying the mutation F193S. However, this clade along with its sister clades has been decreasing in recent month and it doesn’t show a consistent signal of antigenic change. The F193S mutation is observed sporadically within the 3c2.a clade as well.

Additionally, an analysis of “local branching index” [4] suggests that the E62G/R142G/T135K subclade within clade 5 is the most rapidly growing of circulating viruses and is thus estimated to have the highest fitness. With the exception of background 3c2.a1 viruses, other circulating 3c2.a viruses are seen to have similar local branching index values.

**Figure 8.**
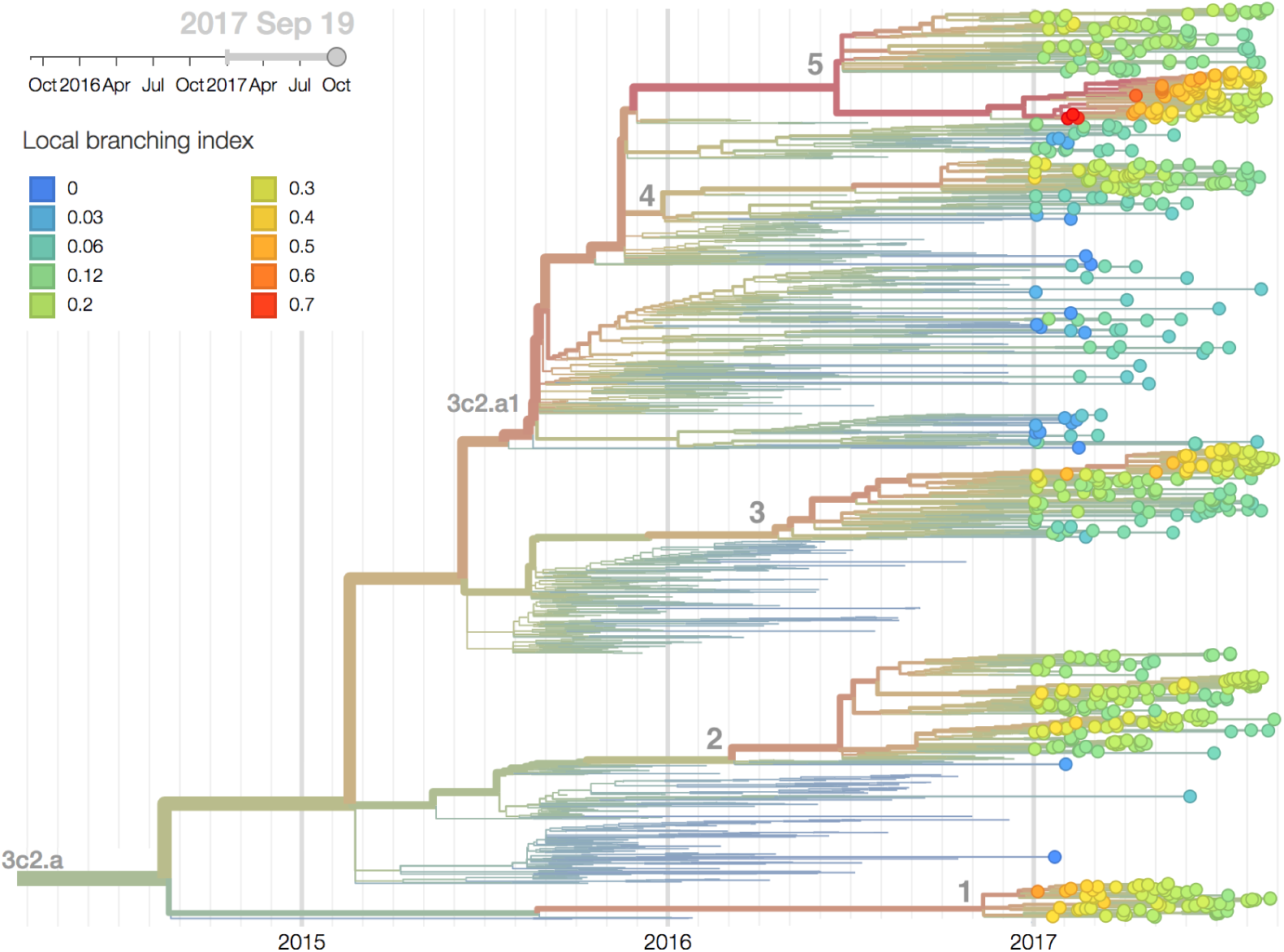
H3N2 / 3c2.a phylogeny colored by local branching index. The tree, zoomed into clade 3c2.a, is colored by local branching index.

*In summary, we think that the clade most likely to increase further in frequency is the E62G/R142G/T135K subclade within clade 5. However, clade 2 viruses (characterized by N121K/R144K) and clade 3 viruses (characterized by T131K/R142K/R261Q) also appear competitive (they are present at moderate frequency, have slightly diverged antigenic phenotypes and are increasing in frequency), although less so than E62G/R142G/T135K viruses (which are growing faster and appear more diverged in antigenic phenotype). We expect all three of these clades to contribute to the future H3N2 population.*

## A/H1N1pdm

A clade comprising mutations S74R and I295V has recently risen to *>*60% global frequency. Although it shows no antigenic distinction by ferret HI data, the rapidity of its rise suggests a selective origin.

We base our primary analysis on a set of viruses collected between Oct 2015 and Aug 2017, comprising approximately 100 viruses per month during relevant months of 2017 (Fig. 9). We use all available data when estimating frequencies of mutations and weight samples appropriately by regional population size and relative sampling intensity to arrive at a putatively unbiased global frequency estimate. Phylogenetic analyses are based on a representative sample of about 2000 viruses.

**Figure 9.**
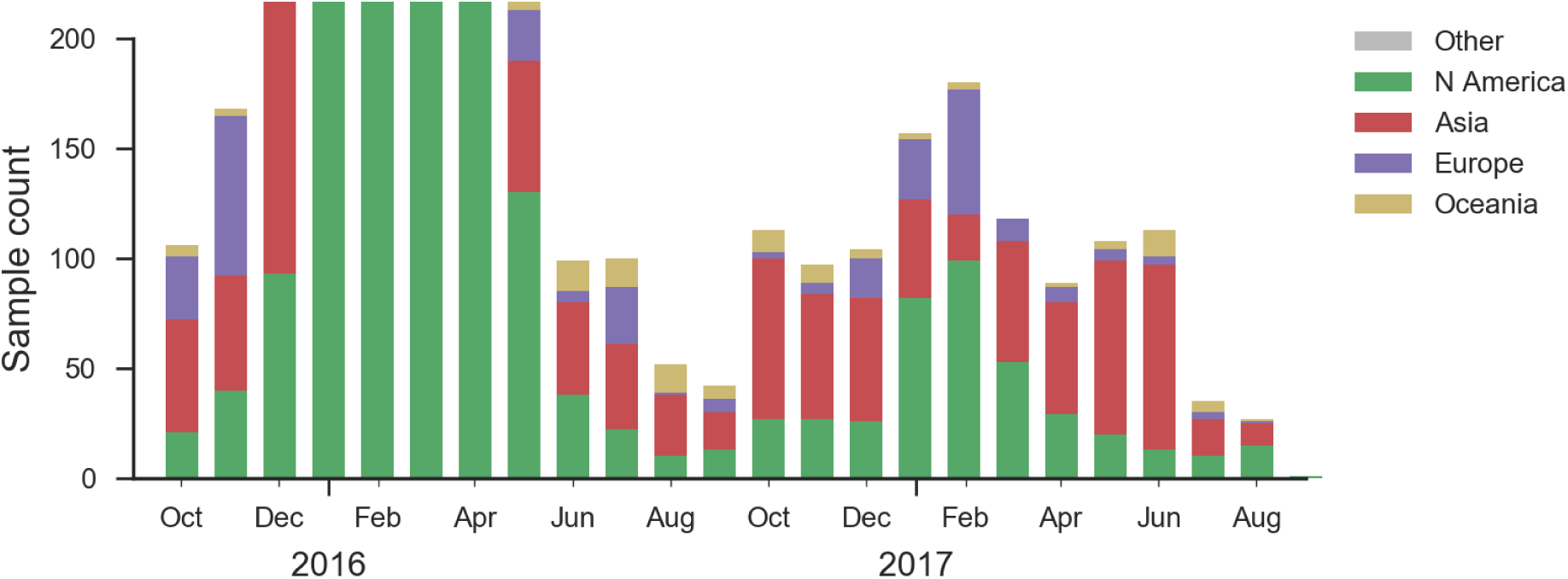
A/H1N1pdm sample counts through time and across regions. This is a stacked bar plot, so that in many months there are more than 100 samples.

H1N1pdm recently underwent a selective sweep of the 6b.1 clade comprising mutations S162N and I216T. Nearly all currently circulating H1N1pdm viruses are clade 6b.1. The H1N1pdm vaccine strain was updated in Sep 2016 to match 6b.1 viruses.

There has been little diversification within clade 6b.1 with a couple notable exceptions. There is a clade comprising mutations S74R and I295V that has been very successful. Within this clade, there is a more recent subclade with the additional mutation S164T.

The S74R and I295V clade has risen rapidly in frequency starting around Jan 2017 (Fig. 10). This clade has been very successful and is now at approximately 66% global frequency. Notably, this clade is at *>*90% frequency in North America, Europe and Oceania but is at lower frequency in Asia. Within this clade the subclade comprising S164T has risen in frequency from April 2017 and is now at approximately 32% global frequency.

**Figure 10.**
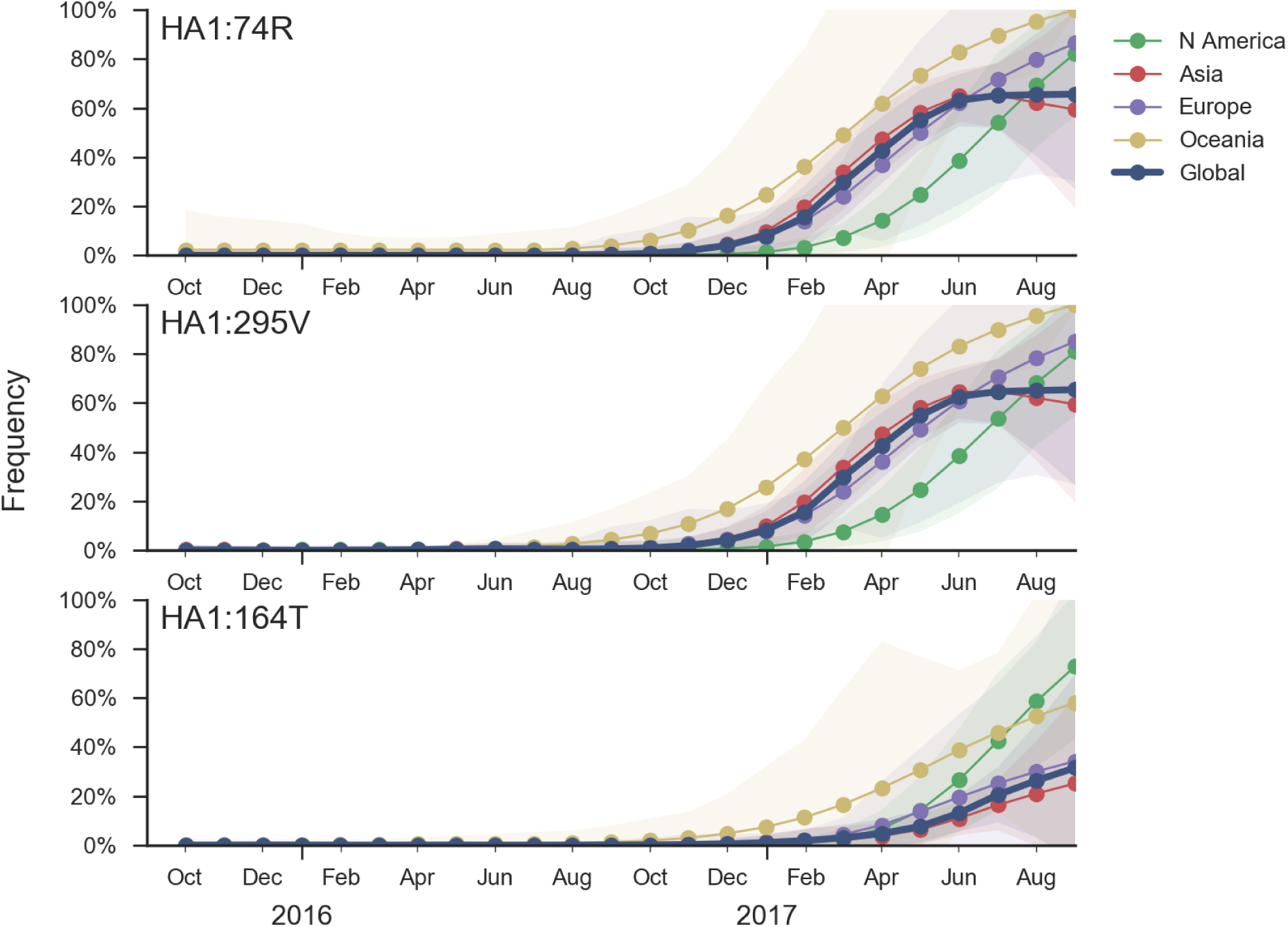
Frequency trajectories of recent mutations in A/H1N1pdm viruses. We estimate frequencies of different mutations based on sample counts and collection dates. These estimates are based on all available data and global frequencies are weighted according to regional population size and relative sampling intensity. We use a Brownian motion process prior to smooth frequencies from month-to-month. Transparent bands show an estimate the 95% confidence interval based on sample counts.

Other mutations have appeared within 6b.1 viruses including S183P, R205K and I324V, but none of these have been very successful and are now declining in response to the rise of S74R and I295V.

Antigenic analysis using HI measurements provided by the Influenza Division at the US CDC fails to show any distinction within recently circulating H1N1pdm viruses (Fig. 11). Like the clades 6b and 6b.1 clade before it, the new S74R and I295V clade does not show any titer drop despite there being ample measurements.

**Figure 11.**
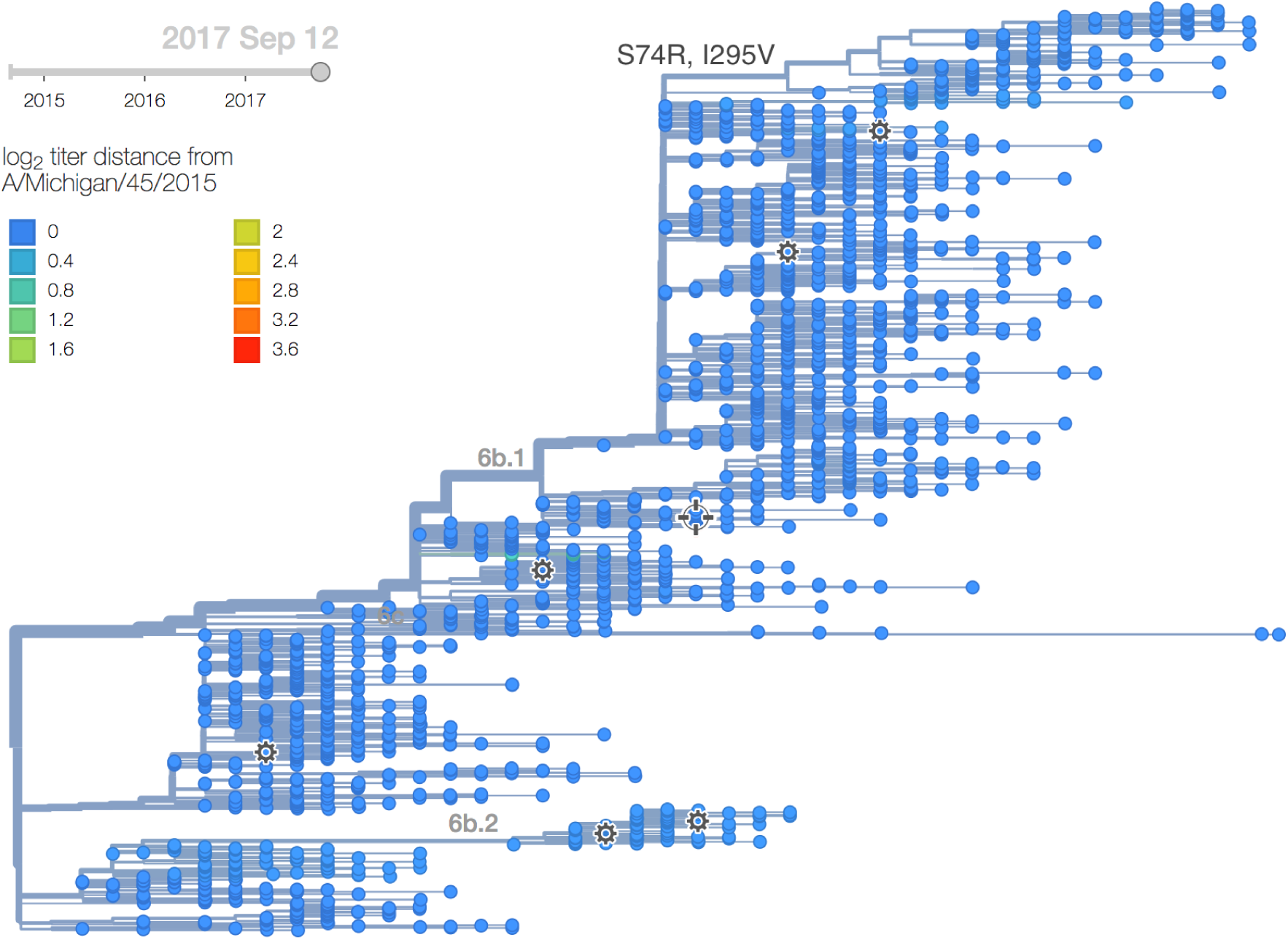
Antigenic distance from A/Michigan/45/2015. Model estimates of antigenic divergence relative to A/Michigan/45/2015. No antigenic variation is picked up by the model.

It was previously observed that clade 6b viruses with mutation K163Q showed decreased titers in a subset of adult human sera due to immunodominance effects, despite continued high titers in ferret sera [5]. Importantly, the new S74R and I295V clade has spread at a similar rate as previous clades 6b and 6b.1 (Fig. 12). This rate of spread and displacement of existing viral diversity is consistent with a selective origin.

**Figure 12.**
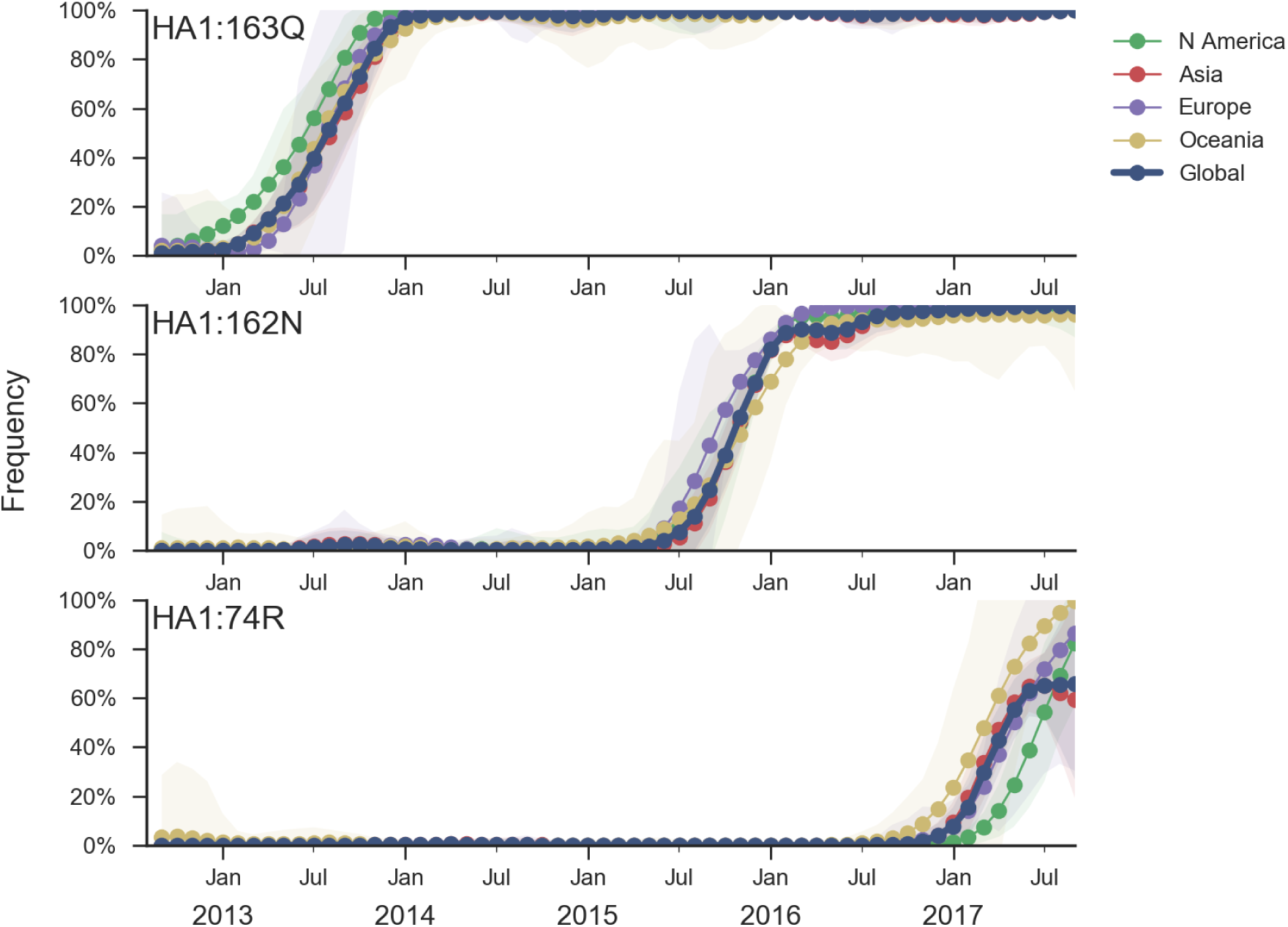
Sequential rapid replacement of clades in A/H1N1pdm. The latest rapid rise of 74R took place at similar or greater speed as the spread of 162N.

*Given current patterns of growth and decline it is highly likely that the S74R and I295V clade will dominant in a years time. Although there is no antigenic effect visible via HI from ferret antisera, the rapidity of the rise of S74R and I295V suggests a selective origin. The S164T subclade also appears to be selectively driven. We would suggest a detailed examination of human serological data as it’s already been established that human serology may differ from ferret serology in H1N1pdm viruses.*

## B/Vic

A clade with a two amino acid deletion 162-/163-has altered serological properties and is increasing in frequency, albeit slowly. Two other clades (carrying mutations K209N and V87A/I175V) have increased in frequency moderately.

We base our primary analysis on a set of viruses collected between Oct 2015 and Aug 2017, comprising between 50 and 250 viruses per month during relevant months of 2017, see Fig. 13. We use all available data when estimating frequencies of mutations and weight samples appropriately by regional population size and relative sampling intensity to arrive at a putatively unbiased global frequency estimate. Phylogenetic analyses are based on a representative sample of about 2000 viruses.

**Figure 13.**
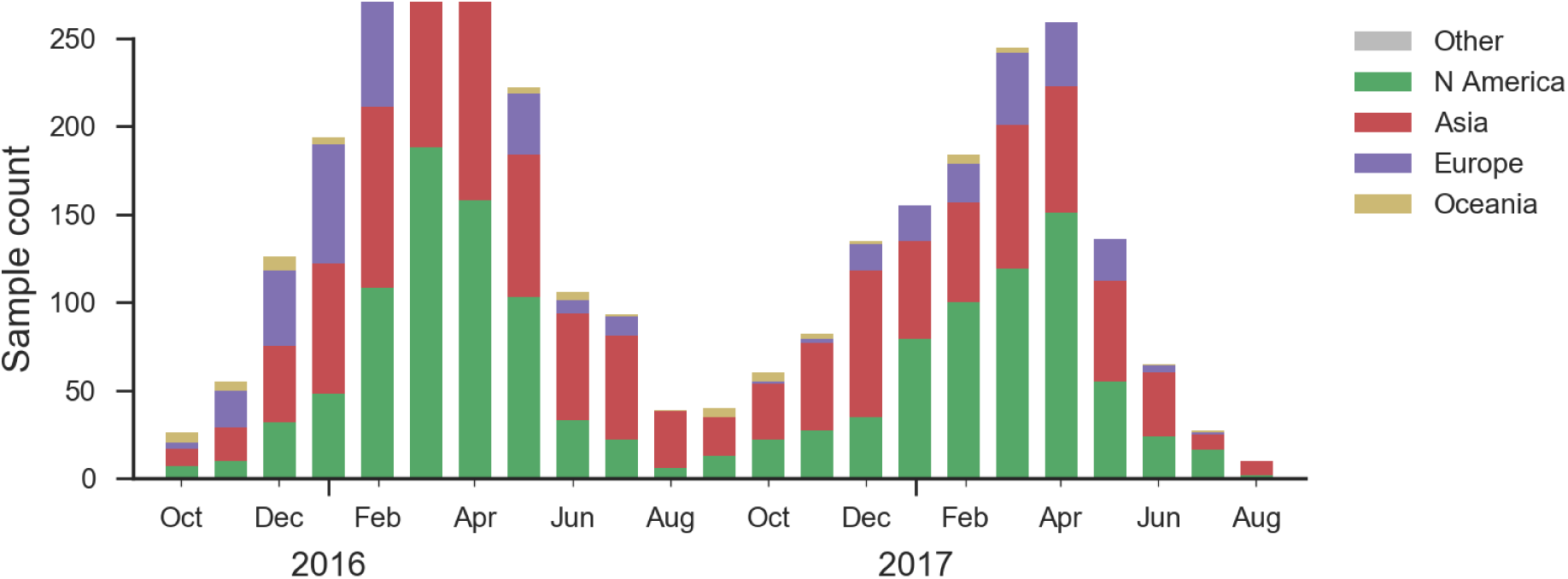
Sample counts through time and across regions. This is a stacked bar plot, so that in good months there are *∼*200 total samples.

**Figure 14.**
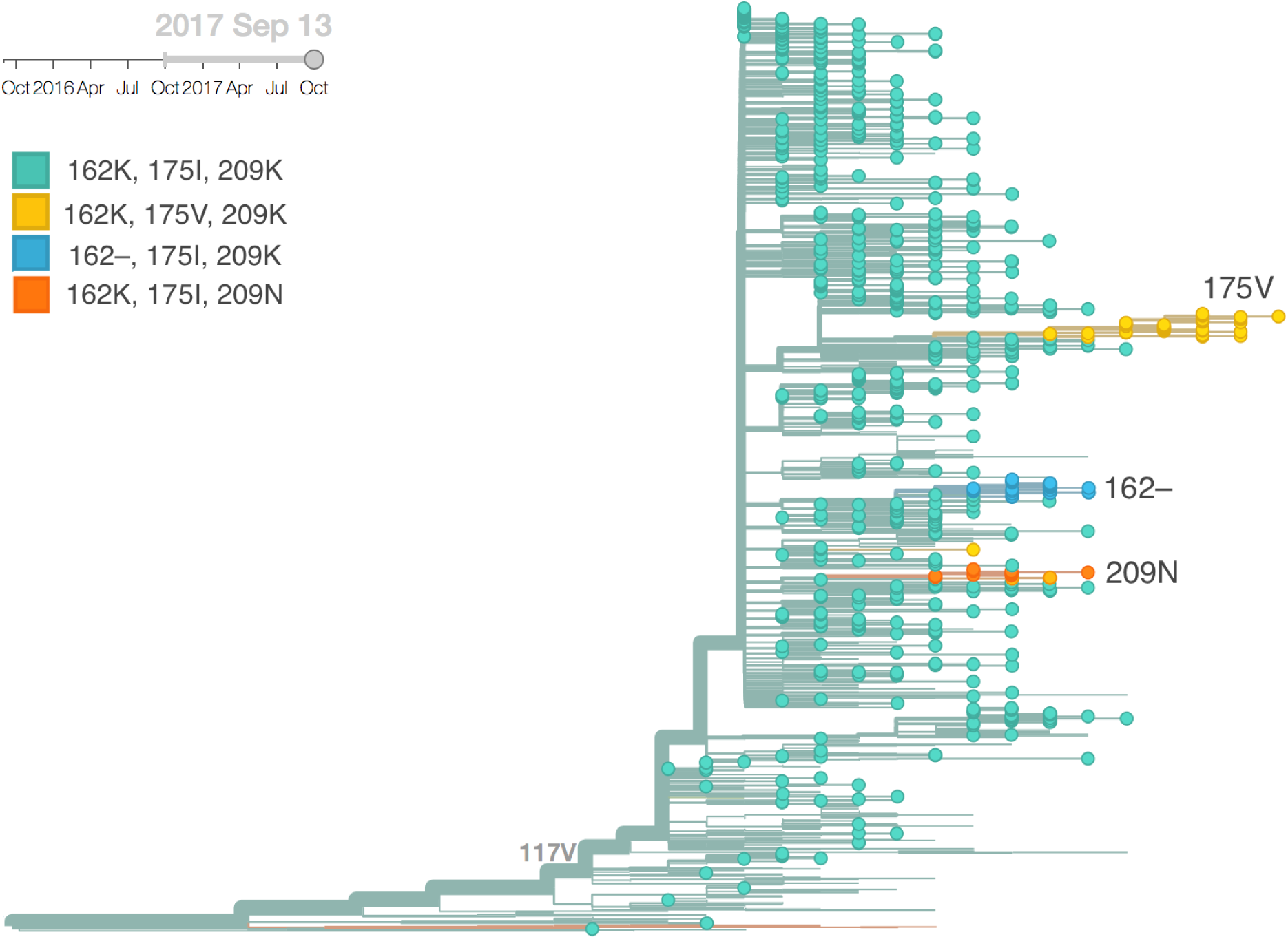
B/Vic phylogeny colored by genotype at positions 162, 175 and 209.

The mutation I117V has continued its slow ascent in frequency as is now *>*99% in the global population. Within this clade there is relatively little circulating genetic diversity (Fig. 14). The only clades of note are a small clade with mutations I180V, D129G and deletions K162- and K163-, another small clade with mutations V87A and I175V and a third small clade with mutation K209N.

The clade bearing 129G, 162–, 163– and 180V has remained at low frequency globally with current global estimates at ∼4%, see Fig. 15. The clade bearing 87A and 175V has increased in frequency substantially in North America but remains low in the rest of the world. The clade bearing 209N has been more successful, approaching 15% globally and circulating in Asia.

**Figure 15.**
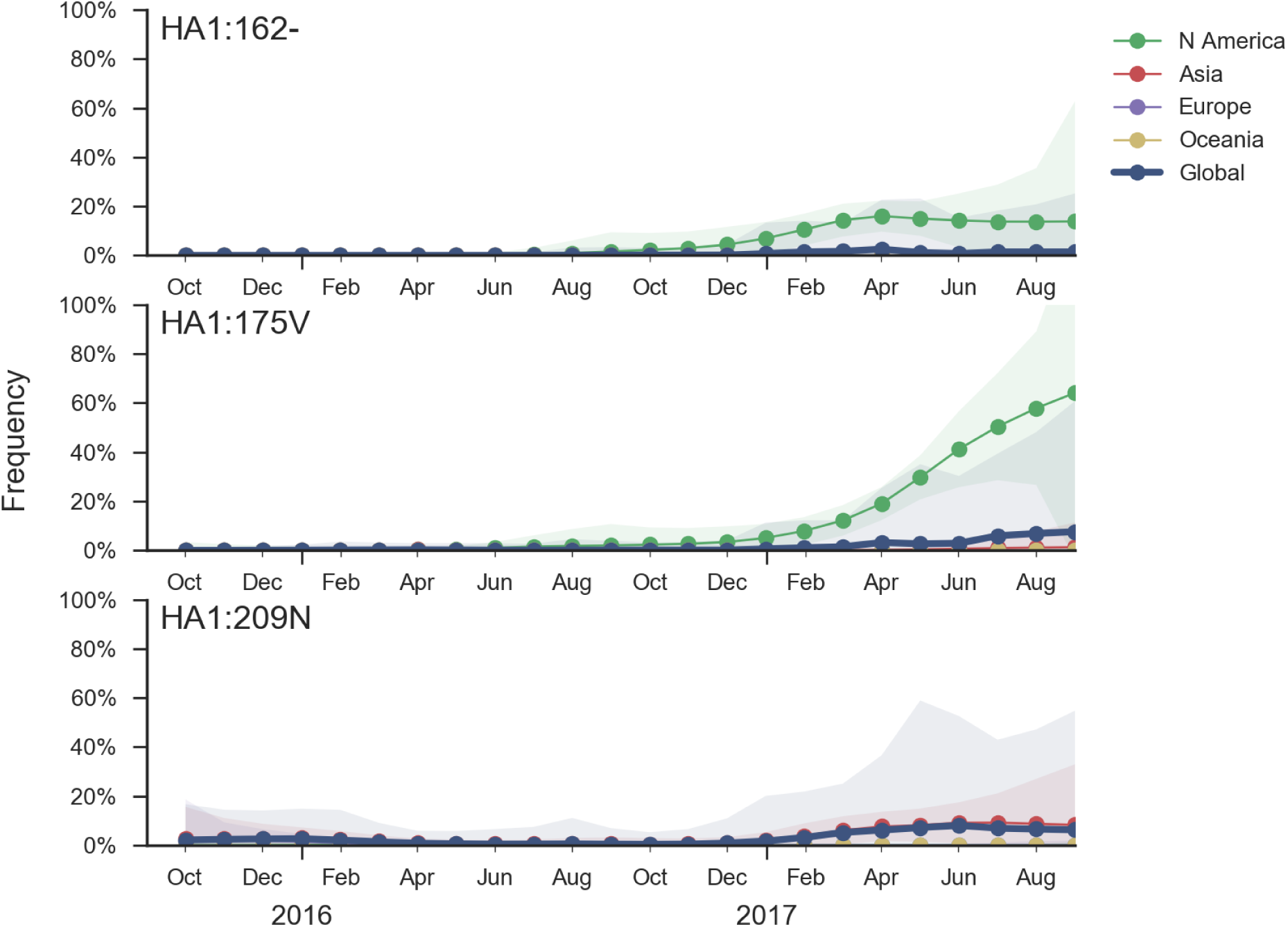
Frequency trajectories of B/Vic variants. We estimate frequencies of different clades based on sample counts and collection dates. We use a Brownian motion process prior to smooth frequencies from month-to-month. Transparent bands show an estimate the 95% confidence interval based on sample counts.

The clade bearing 129G, 162–, 163– and 180V appears clearly drifted according to HI measurements provided by the Influenza Division at the US CDC (Fig. 16). We estimate an approximately 8-fold titer drop (3 antigenic units) of drift from vaccine strain B/Brisbane/60/2008. Separately, a small number of recent viruses sampled in Hong Kong have a 9bp deletion overlapping codons 162, 163, 164, and 165 along with a mutation I180T. While based on limited data, the tree model estimates an 1.2 unit titer drop.

**Figure 16.**
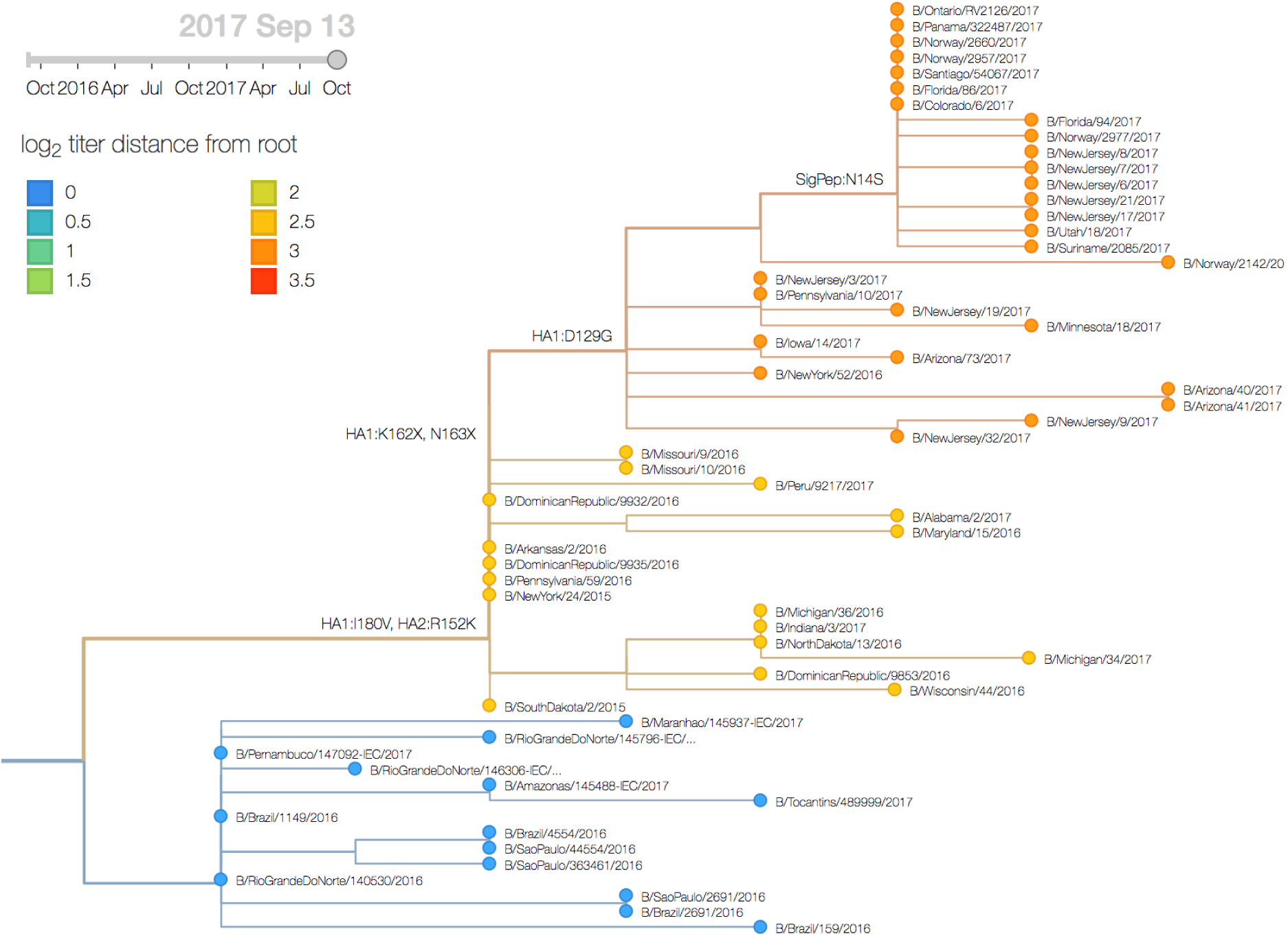
Antigenic evolution of the B/Vic 162-/163-clade. The tree model estimates a drop in HI titers by 3 units.

*Although the clade bearing 129G, 162–, 163– and 180V is almost certainly antigenically drifted, its slow spread up to the this point suggests that it is unlikely to predominate in a year’s time. We would advise careful monitoring as further mutations may enhance spread or similar mutations might emerge on fitter backgrounds.*

## B/Yam

A clade comprising M251V within clade 3 viruses continues to dominate. The is little genetic differentiation within this clade and no evidence of antigenic evolution.

We base our primary analysis on a set of viruses collected between Oct 2015 and Aug 2017, comprising between 50 and 250 viruses per month during relevant months of 2017 (Fig. 17). We use all available data when estimating frequencies of mutations and weight samples appropriately by regional population size and relative sampling intensity to arrive at a putatively unbiased global frequency estimate. Phylogenetic analyses are based on a representative sample of about 2000 viruses.

**Figure 17.**
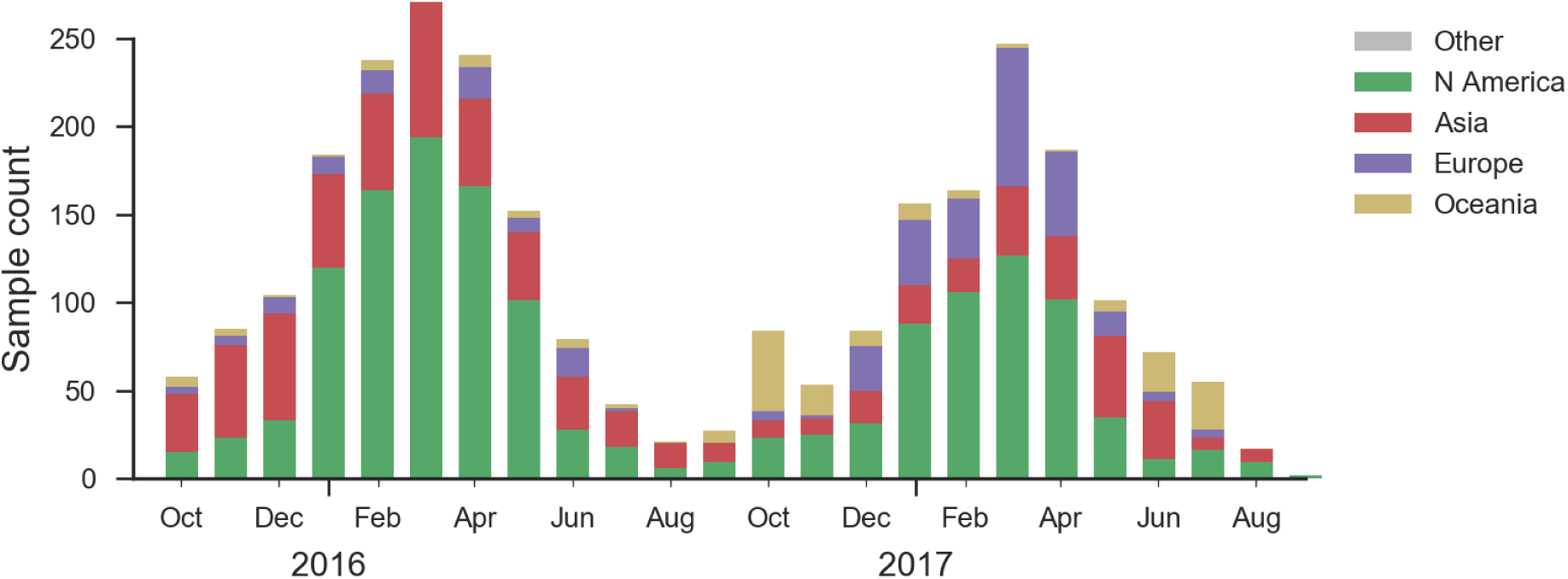
Sample counts through time and across regions. This is a stacked bar plot, so that in good months there are *∼*200 total samples.

Clade 2 viruses appear to be extinct from the world. Clade 3 viruses have continued to evolve and genetic diversity relative to vaccine strain B/Phuket/3073/2013 has emerged. Of note the clade comprising M251V dominates the population with over 99% frequency (Fig. 18). The subclade within M251V comprising K211R has decreased in frequency and is now extinct or nearly extinct. The only notable subclade within the M251V clade is that bearing T76I which is circulating at low frequency.

**Figure 18.**
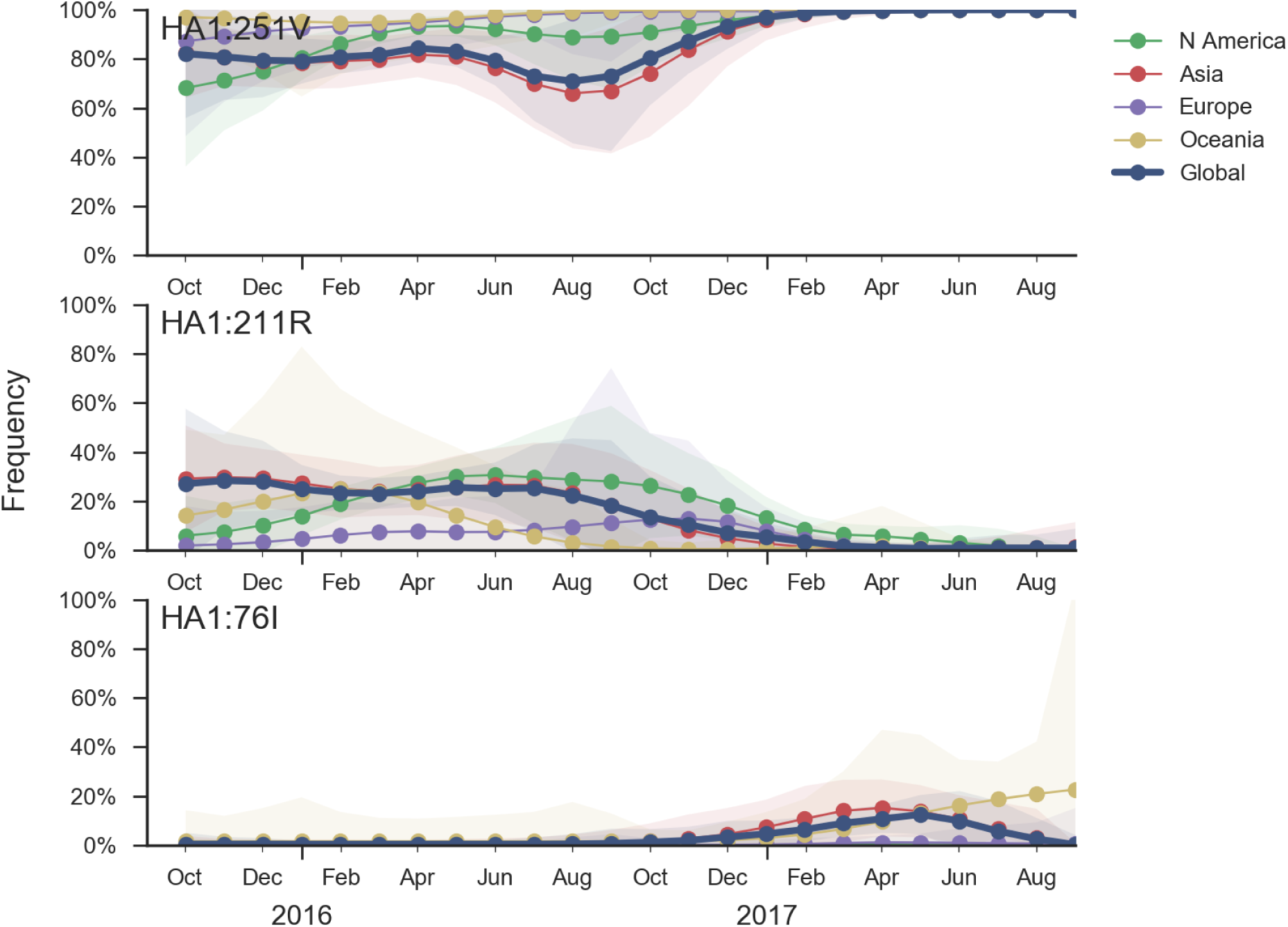
Frequency trajectories of B/Yam variants. We estimate frequencies of different clades based on sample counts and collection dates. We use a Brownian motion process prior to smooth frequencies from month-to-month. Transparent bands show an estimate the 95% confidence interval based on sample counts.

Based on HI measurements provided by the Influenza Division at the US CDC, we find no evidence for antigenic evolution within clade 3 viruses and the vaccine B/Phuket/3073/2013 remains well matched to circulating viruses (Fig. 19).

**Figure 19.**
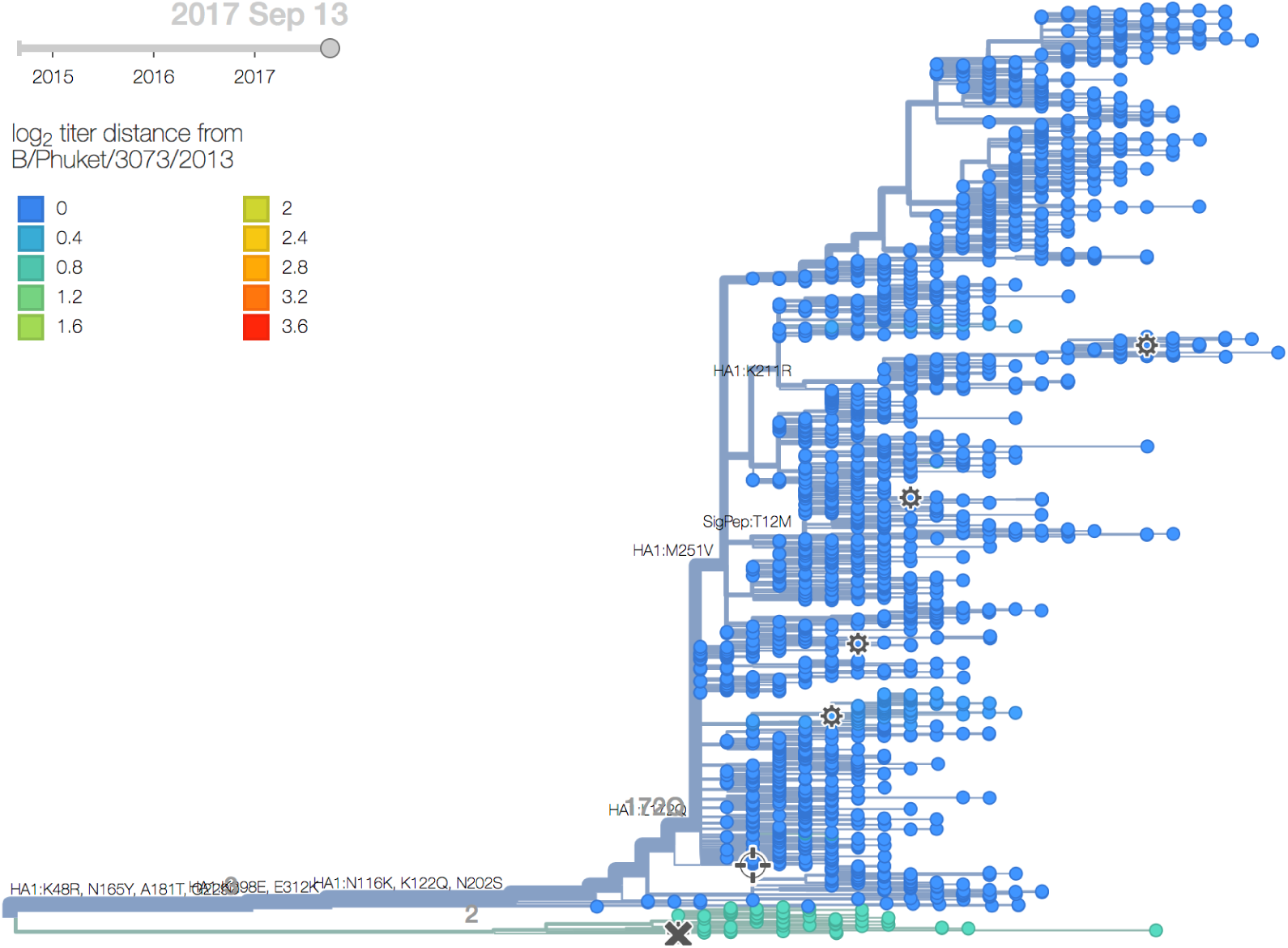
B/Yam phylogeny colored by antigenic distance from B/Phuket/3073/2013. The tree model of antigenic distances infers no antigenic evolution.

*Barring the emergence of a new selective variant we anticipate relative stasis of B/Yam viruses in the coming year.*

